# Enhanced Specificity Mutations Perturb Allosteric Signaling in CRISPR-Cas9

**DOI:** 10.1101/2021.09.14.460226

**Authors:** Łukasz Nierzwicki, Kyle W. East, Uriel N. Morzan, Pablo R. Arantes, Victor S. Batista, George P. Lisi, Giulia Palermo

## Abstract

CRISPR-Cas9 is a molecular tool with transformative genome editing capabilities. At the molecular level, an intricate allosteric signaling is critical for DNA cleavage, but its role in the specificity enhancement of the Cas9 endonuclease is poorly understood. Here, solution NMR is combined with multi-microsecond molecular dynamics and graph theory-derived models to probe the allosteric role of key enhancement specificity mutations. We show that the mutations responsible for increasing the specificity of Cas9 alter the allosteric structure of the catalytic HNH domain, impacting the signal transmission from the DNA recognition region to the catalytic sites for cleavage. Specifically, the K855A mutation strongly disrupts the HNH domain allosteric structure, exerting the highest perturbation on the signaling transfer, while K810A and K848A result in more moderate effects on the allosteric intercommunication. This differential perturbation of the allosteric signaling reflects the different capabilities of the single mutants to increase Cas9 specificity, with the mutation achieving the highest specificity also strongly perturbing the signaling transfer. These outcomes reveal that the allosteric regulation is critical for the specificity enhancement of the Cas9 enzyme, and are valuable to harness the signaling network to improve the system’s specificity.

## INTRODUCTION

CRISPR-Cas9 (clustered regularly interspaced short palindromic repeat and associated Cas9 protein) is a bacterial adaptive immune system with widely demonstrated and profound genome editing capabilities.^1^ At the core of the CRISPR-Cas9 technology, the endonuclease Cas9 can be programmed with single guide RNAs to site-specifically recognize and cleave any desired DNA sequence, enabling easy manipulation of the genome and playing a pivotal role in gene editing applications.^2^ The RNA-programmable Cas9 generates double stranded breaks in DNA by first binding complementary DNA sequences and then using two endonuclease domains, HNH and RuvC. Structures of the *Streptococcus Pyogenes* Cas9 (SpCas9) revealed that a large recognition lobe (REC) mediates the nucleic acid binding through three regions (REC1-3), while the HNH and RuvC nucleases are separated and act as molecular scissors of the two DNA strands (Fig. 1).^3^

**Figure 1.**
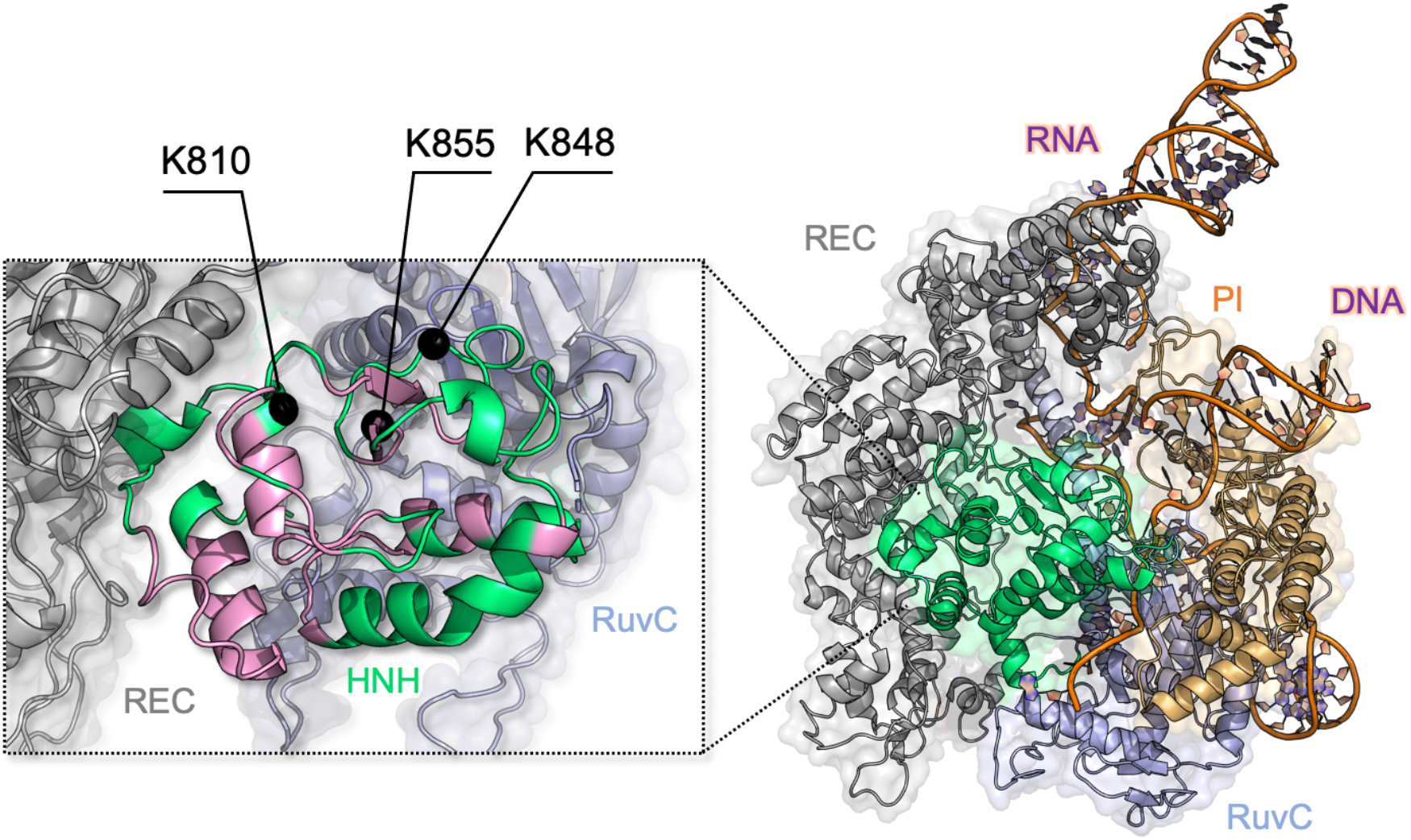
Architecture of the Cas9 endonuclease (PDB: 5F9R)^9^ highlighting its protein domains as follows: HNH (green) RuvC (light blue), PAM-interacting (PI, orange) and recognition lobe REC (gray). A portion of the RNA:DNA hybrid behind HNH has been removed for clarity. Close-up view (left): the allosteric pathway spanning the HNH domain is shown (pink) and the locations of the three specificity-enhancement K–*to*–A mutation sites are labeled.^11^

Biophysical studies revealed that the molecular function of CRISPR-Cas9 is characterized by an intricate allosteric communication, which is critical for transferring the information of DNA binding from the REC lobe to the catalytic sites for cleavage.^4–8^ Biochemical and single-molecule experiments suggested that the catalytic HNH domain is the core of this allosteric relay.^4,5^ Indeed, the high flexibility of HNH can facilitate the signal transduction,^9,10^ exerting a conformational control over double stranded DNA cleavage.^4^ Solution NMR and all-atom molecular dynamics (MD) have indicated a dynamic pathway of allosteric residues through HNH, depicting a mechanism for biological information transfer.^11^ The identified contiguous dynamic network crosses the HNH domain, propagating the DNA binding signal from the REC region to the nuclease domains (HNH and RuvC, Fig. 1, left panel) for concerted cleavages of the two DNA strands.^4^ This provided a route for the allosteric transduction, and a mechanistic rationale to single-molecule and biochemical experiments,^4,5^ clarifying how the HNH dynamics could transfer the information of DNA binding from the REC region to the cleavage sites. This REC-HNH-RuvC allosteric communication is also critical for the system’s specificity. Indeed, the binding of off-target DNA sequences at the level of the REC lobe results in alterations of the dynamics of HNH and, in turn, affects the DNA cleavage capability of Cas9.^5,6,12^ To improve the system’s specificity and reduce its off-target activity, extensive engineering of the Cas9 protein has been performed.^5,13–15^ Three lysine-to-alanine (K–*to*–A) point mutations (i.e., K810A, K848A and K855A) within the HNH domain have shown to be important for the specificity enhancement,^14^ but their mode of action has remained unmet. This knowledge is of major importance, as it could help the mechanism-based design of improved Cas9 variants.

Here, we probe the structural and dynamic role of these mutations with respect to the HNH allosteric signaling via solution NMR, molecular simulations and network models derived from graph theory. This integrative approach holds the capability of defining allosteric motions with experimental accuracy through NMR^16–19^ while also describing the network of communication at atomic level detail through computational methods.^20–27^ We reveal that the three enhanced specificity mutations alter the HNH allosteric structure, impacting the signal transmission from REC to RuvC. Indeed, the K855A mutation strongly disrupts the communication mediated by the HNH allosteric core, exerting the highest perturbation on the REC-HNH-RuvC allosteric communication. K810A and K848A result in more moderate effects on the allosteric intercommunication. This differential perturbation of the allosteric signaling reflects the different capability of the three single mutants to increase Cas9 specificity,^14^ suggesting that alterations of the allosteric signaling could be critical for the specificity enhancement. Taken together, our findings disclose that enhanced specificity mutations perturb the HNH allosterism, which in turn impacts Cas9 specificity. These findings represent a decisive step forward in understanding of role of allostery in the specificity of Cas9, and contribute engineering efforts toward improved genome-editing.

## RESULTS

Solution NMR has been used to trace allosteric motions within a construct of the HNH domain that shows consistency with the structure of HNH within the full-length CRISPR-Cas9 system (Fig. 2A).^11^ Indeed, due to the size of its polypeptide chain (i.e., 160 kDa), the Cas9 protein challenges traditional solution NMR, requiring optimized constructs of the individual domains to report on its structural and dynamical features.^11,28^ To address the allosteric mechanism, it is however critical to characterize the communication network within the full-length Cas9. Toward this aim, we performed all-atom MD simulations of the CRISPR-Cas9 system in its wild-type (WT) form^9^ and introducing the K848A, K810A and K855A mutations. These full-length systems, comprising >340,000 atoms each, have been simulated reaching μs trajectory length and in replicas, collecting a solid multi-μs statistic al ensemble for the analysis of the allosteric mechanism (details are reported in the Methods section).^7,11^ Molecular simulations also considered the isolated HNH domain,^11^ to assess the structure and dynamics of HNH in its isolated form.

**Figure 2.**
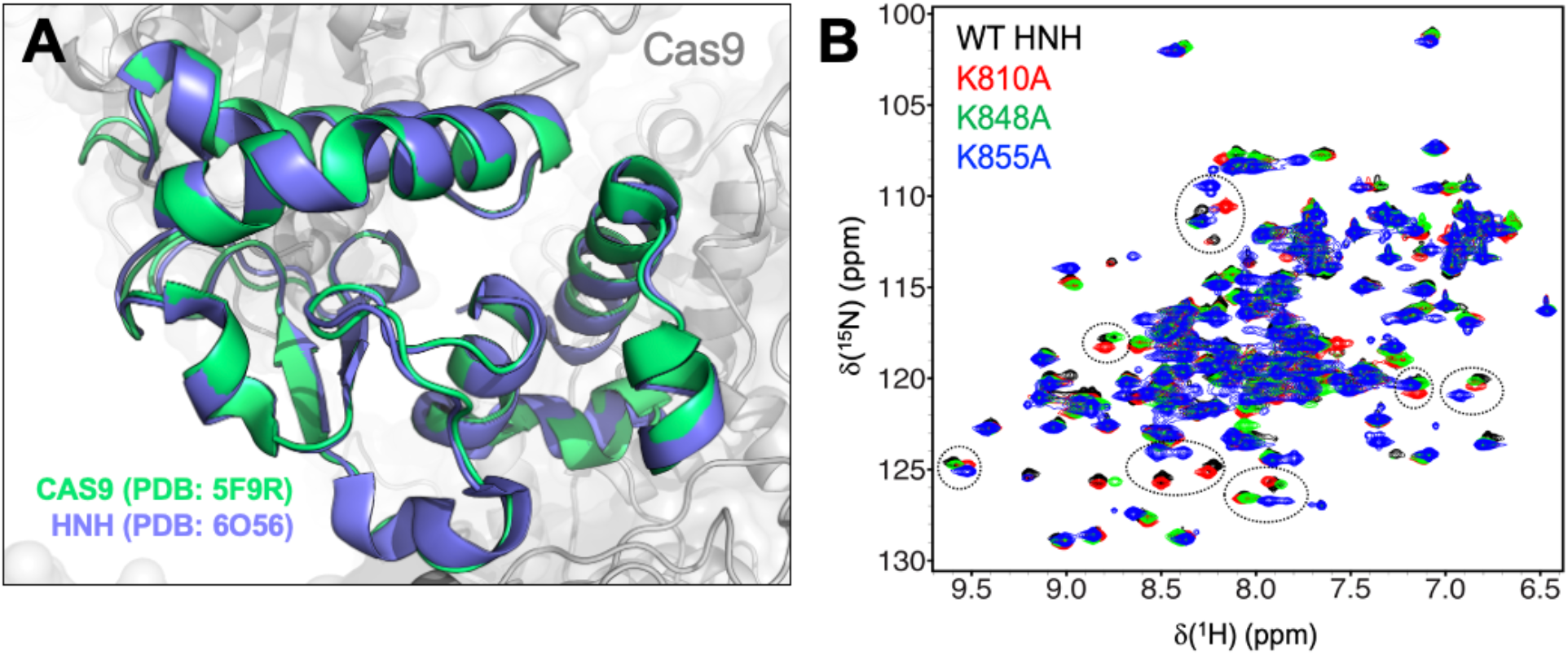
**(A)** The structure of the HNH domain (residues 775-908) crystallized within the full-length Cas9 (PDB: 5F9R, green)^9^ is superposed on the X-ray structure of the HNH construct (PDB: 6O56, blue).^11^ The two structures display an RMSD difference for the Cα below 1 Å (i.e., 0.92 Å). **(B)** NMR spectra of the wild-type (WT) HNH domain and its mutants. The ^1^H-^15^N HSQC NMR spectra of K810A (red), K848A (green), and K855A (blue) are overlaid with that of WT HNH (black). Selected areas of strong chemical shift perturbation are circled on the spectra.

### Structural perturbation of the HNH endonuclease

We used targeted mutagenesis to introduce the three previously identified specificity enhancement K–*to*–A mutations (K810A, K848A and K855A)^14^ into the HNH construct,^11^ and we employed solution NMR to determine the structural changes associated with these point mutations (Fig. 2). First, changes to the local structure of HNH caused by each mutation were derived from chemical shift (Δδ) perturbations in ^1^H-^15^N HSQC NMR spectra (Fig. 2B). Despite significant chemical shift perturbations are observed, the overall structure of HNH is maintained in the mutants, as also confirmed by circular dichroism analysis revealing that all systems are > 95 % folded at the temperature of the experiments (25 °C, Fig. S1). The environmental perturbations were calculated using the method of Bax and coworkers.^29^ For each mutant, the composite Δδ ^1^H-^15^N, as well as the total Δδ perturbations are reported in Fig. 3. Based on NMR measurements, it is apparent that each point mutation has a unique effect on the HNH domain. In detail, K855A displays severe exchange broadening (where the signal-to-noise has decreased by over 20-fold, Fig. 3A, gray vertical bars) throughout the core of HNH (residues 842, 848, 849, 851, and 858, Fig. 3B) and many significant Δδ perturbations on the RuvC-adjacent interface (residues 780, 813, 821-828, 838-841, 850, 853, 856-872, and 903-908). These regions are integral part of the HNH allosteric pathway previously identified (Fig. 3B).^11^ K810A decreases the Δδ perturbation (residues 812-813, 834, 840-846, and 856), while K848A causes moderate chemical shifts effects (residues 849 and 896). Most notably, the overall degree of Δδ perturbation decreases from K855A > K810A > K848A (Fig. 3C, black bars). Intriguingly, this trend in the structural perturbation mirrors the location of these residues with respect to the WT HNH allosteric pathway (Figs. 1 and 3B). Indeed, K855A, which is fully embedded in the allosteric pathway, shows the largest Δδ perturbation. K810A, being on the periphery of the pathway, decreases the Δδ perturbation, while K848A, being distant from the pathway, causes more moderate chemical shift effects. These observations suggest that K855A could have a more pronounced effect the HNH allosteric structure, while K810A and K848A could exert a more moderate effect. This observation and its implications on the HNH allosterism are thoroughly analyzed in the forthcoming (*vide infra*).

**Figure 3.**
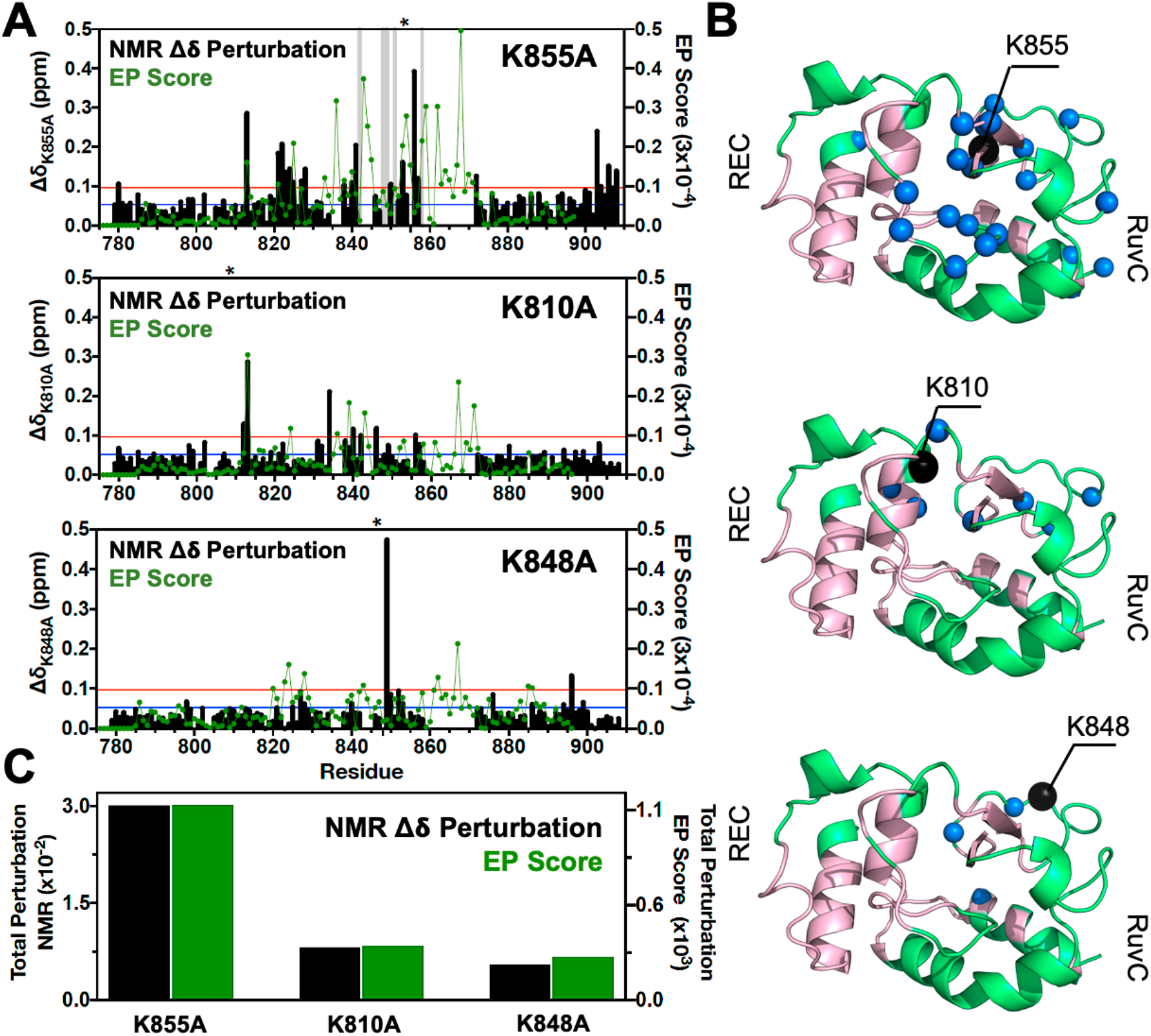
Environmental perturbations caused by the K855A, K810A and K848A mutations in HNH. **(A)** The NMR composite chemical shifts (black bars) determined by 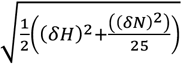 are reported for each mutant. Blue lines are the 10% trimmed mean of all shifts and red lines represent 1.5σ above the 10% trimmed mean. Sites of severe line broadening are represented by light gray bars. The site of mutation is indicated using an asterisk (*) above each plot. The Environmental Perturbation (EP) scores computed for the isolated HNH domain from MD simulations are shown using a green line. Blue and red lines from chemical shift analysis also represent the mean (+ 1σ) of the EP score data (right y-axis). **(B)** Residues with composite chemical shifts above the significance cutoff (blue spheres) are plotted onto the crystal structure of the HNH domain (green). The WT HNH allosteric pathway is also shown (pink). (**C**) Total chemical shift and environmental perturbation (black and green bars respectively) obtained as the sum of the NMR chemical shift perturbations and of the EP scores respectively, for the isolated HNH domain.

### Perturbation of the chemical environment

To determine the structural perturbation induced by the three K–*to*–A mutations from MD simulations, we introduced an Environment Perturbation (EP) score, which determines the extent of the dynamic perturbation for a specific atom, given its local environment (details are in the Methods section). As a result, the EP scores computed for the isolated HNH domain peak around amino acid positions similar to the NMR Δδ of the HNH construct (Fig. 3A). This shows a qualitative agreement between the NMR and MD environmental perturbations, which is reasonable considering that the NMR Δδ are direct reporters of the local environment. The 860-870 region remained unassigned in the NMR spectra, likely due to the remarkable flexibility of the region, which in turn results in high computed EP scores. It is also notable that the environmental perturbations computed for the HNH domain within the full-length Cas9 (Supplementary Fig. S2) are consistent with the EP scores of the HNH domain in its isolated form.

Overall, we also observe that the total environmental perturbation arising from MD simulations follows the trend of the total NMR Δδ perturbations, since they are both higher in K855A and decreases in K810A and K848A (Fig. 3C). The qualitative agreement between the NMR Δδ perturbations and the computed EP scores indicates that the structural ensemble obtained through all-atom MD for both the full-length Cas9 and the isolated HNH represents well the local environment determined through the NMR Δδ. Ultimately, the computational characterization of the chemical environment follows the experimental trend of the Δδ perturbations (Fig. 3), revealing that the structural perturbation is higher in K855A and decreases in K810A and K848A. The computed EP scores thereby support the idea that K855A could most strongly alter the allosteric structure of HNH, while K810A and K848A could progressively exert a more moderate impact.

### Alteration of the allosteric communication

Here, we combine Carr-Purcell-Meiboom-Gill (CPMG) relaxation dispersion NMR experiments with information theory and community network approaches to shed light on whether the three specificity-enhancement mutations (i.e., K810A, K848A and K855A) could intervene in the HNH-mediated allosteric communication. CPMG relaxation dispersion detect dynamical motions over slow timescales (μs to ms),^30^ which are an indication of the allosteric signaling transfer, as shown in a number of previous studies of allosteric enzymes.^17,31^ CPMG analysis on the three variants detects μs-ms motions in numerous residues comprising the WT pathway (e.g., G790, L791, I795, I841, F846, V856, S872, E873, I892, Q894, L900, Fig. 4 and Supplementary Fig. S3), with slight differences occurring in each of the variants (Supplementary Table S1). This overall consistency confirms the retention of the dynamics of the residues comprising the allosteric pathway in the WT Cas9.^11^ Indeed, the trend of curved dispersion profiles is consistent between the WT and the variants, suggesting that many of the residues in WT HNH remain flexible in the mutants with only minor variations in the amplitude of the curves. These amplitude changes correspond to small ΔR_ex_ ≤ 1.5, which indicates relatively small dynamic changes.

**Figure 4.**
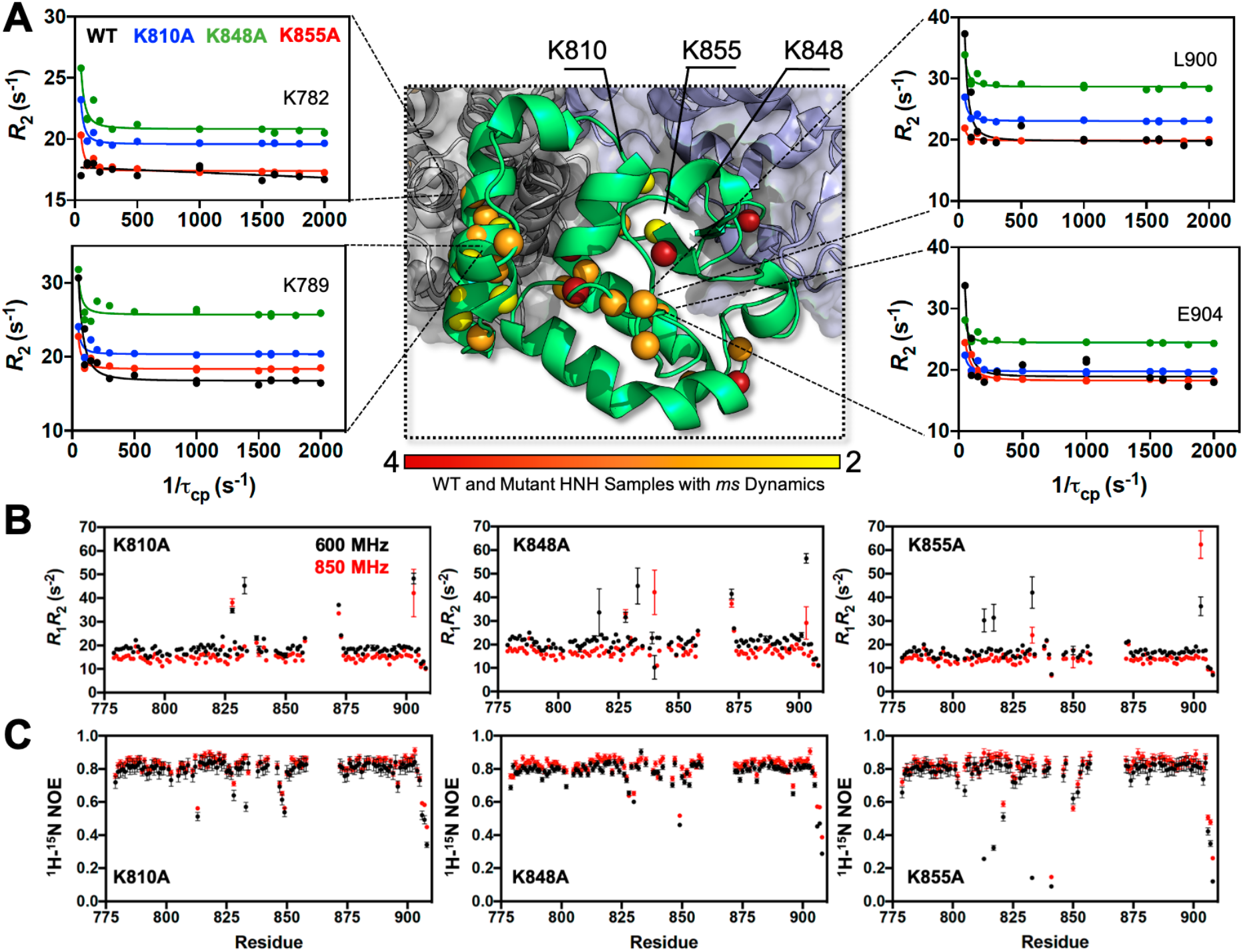
Dynamic properties of HNH and its mutants. (**A**) Structure of the HNH domain showing *ms* timescale dynamics preserved upon mutation of the three K–*to*–A mutations (central panel). Spheres represent sites of CPMG relaxation dispersion that appear in the WT HNH and *at least one other* specificity enhancement variant. These sites preserve the dynamics upon mutations and are color-coded from red (highly preserved dynamics) to yellow (moderately, yet still preserved dynamics). Close-up views of representative CPMG relaxation dispersion profiles for WT (black), K810A (blue), K848A (green), and K855A (red) HNH are shown for various residues in this cluster. (**B**) Plots of the relaxation rate product, *R1R2* for HNH mutants collected at 600 (black) and 850 (red) MHz. (**C**) Plots of the ^1^H-[^15^N] heteronuclear NOE for HNH mutants collected at 600 (black) and 850 (red) MHz.

To describe the allosteric pathway of communication, information theory has been applied to the analysis of our μs-length simulations of the Cas9 mutants. We computed the dynamical pathways composed by residue-to-residue steps that optimize the momentum transport (and thereby maximize the correlations) from REC to RuvC, in analogy the WT system (details are reported in the Methods Section).^11^ We found that the residues composing the dynamical pathway in the three variants differ very little from the WT, and are consistent for the full-length CRISPR-Cas9 and the HNH construct (Supplementary Figs. S4-S5). This indicates that the allosteric dynamic signaling is preserved in the Cas9 mutants, in agreement with CPMG experiments. In this respect, the pathways that maximize the dynamic transmission between RuvC and REC2 through HNH agree well with the slow residues identified in the HNH construct via CPMG relaxation dispersion (Supplementary Figs. S3-S5).

To further understand the effect of the K810A, K848A and K855A mutations on the allosteric mechanism, we employed community network analysis^26^ to identify the groups of residues that comprise cohesive structural units with synchronized dynamics within HNH. These “communities” of highly correlated residues establish a dynamic “cross-talk” with each other, whose strength can be quantified using the “edge betweenness” (EB) measure (details are reported in the Methods section). This analysis has been performed on the full-length CRISPR-Cas9 systems (reported here), and for comparison in the isolated HNH domain (reported in the Supplementary Data). First, community network analysis has been performed on the WT HNH, identifying seven communities (Fig. 5A). Three communities hold most of the residues that display slow dynamics through CPMG relaxation dispersion, and that compose the allosteric pathway in the WT Cas9.^11^ These “allosteric” communities (A1, yellow, A2 cyan and A3 purple) are full part of the allosteric route communicating the RuvC and REC2 interfaces (at the A1 and A3 communities, respectively). The “non-allosteric” communities (NA1 orange, NA2 tan, NA3 red and NA4 black) include only few allosteric residues. To understand how each mutation affects the inter-community cross-talk and the allosteric network of communication, we computed the mutation-induced EB change (*Δ*EB, details are in the Methods section and in the Supplementary Data), allowing us to quantify the perturbation in the communication induced by the mutation. The *Δ*EB have been normalized and plotted using circular graphs (Fig. 5B), where the communities are disposed in a circle and connected using links, whose thickness is proportional to the *Δ*EB. Negative values of *Δ*EB (red) represent loss of communication, as opposed to positive values (blue), which indicate a communication gain upon mutation. For all mutants, we observe a dramatic loss of communication between the A1 and A2 allosteric communities, which are central in the allosteric pathway (Fig. 5B). This loss of communication at the core of the HNH allosteric structure indicates that the substitutions that enhance the specificity of Cas9 disrupt the allosteric cross-talk between RuvC and REC2. This evidence links the enhancement of specificity to the disruption of the allosteric pathway, pinpointing that an increase in specificity induced by the K–*to*–A mutations is associated to alterations of the HNH allosteric signaling.

**Figure 5.**
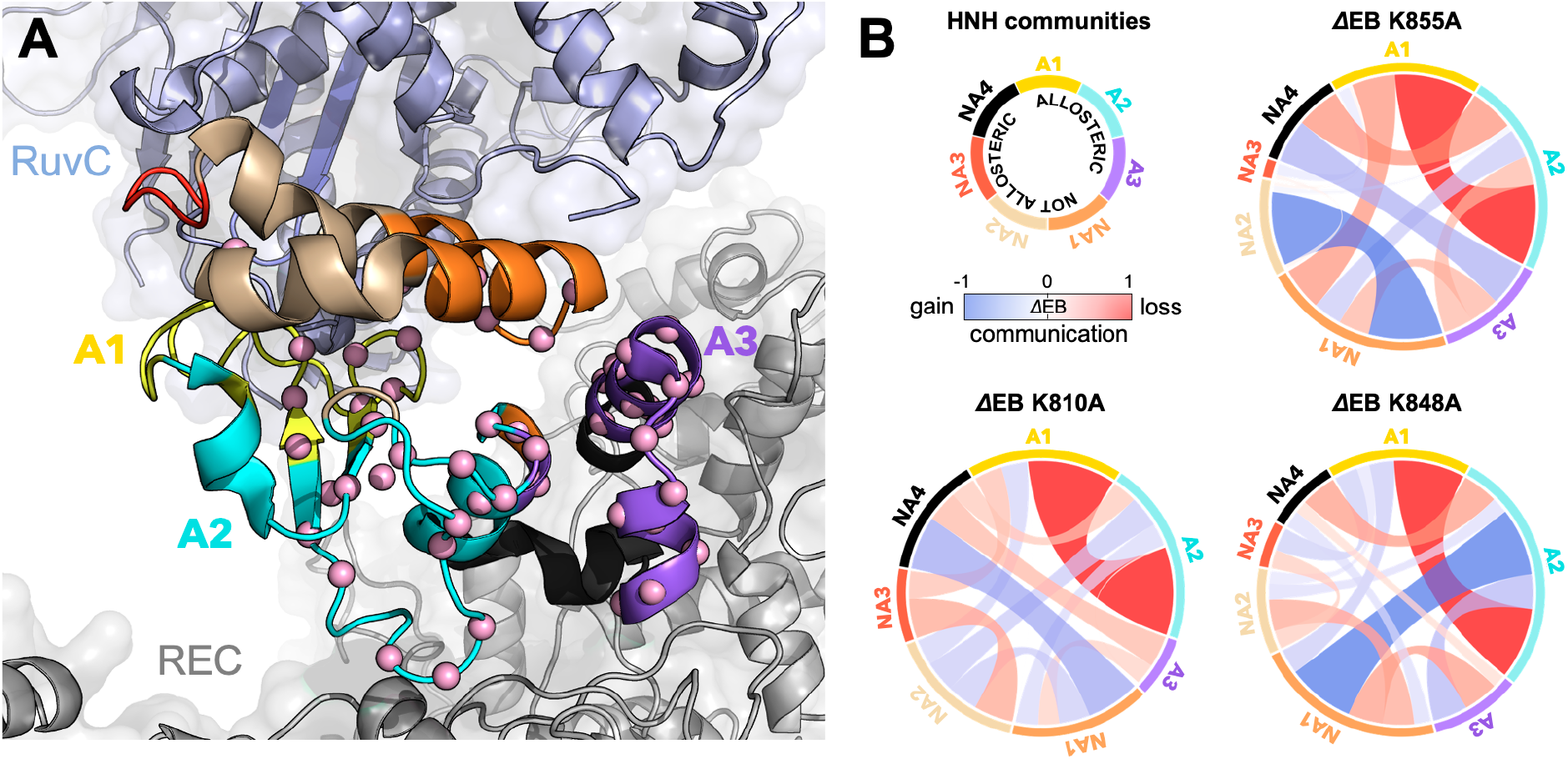
Alterations of the allosteric signaling in the full-length CRISPR-Cas9 systems. **(A)** Close-up view of the HNH domain within the WT full-length Cas9, showing seven communities of synchronized dynamics, indicated using different colors. Three communities are allosteric (A1 yellow, A2 cyan and A3 purple), since they hold most of the residues that compose the allosteric pathway (shown as spheres). The non-allosteric communities (NA1 orange, NA2 tan, NA3 red, NA4 black) include a few allosteric residues. (**B**) Circular graphs reporting the mutation-induced Edge Betweenness change (*Δ*EB), a measure of the communication gain or loss between couples of communities upon mutation. For each of the K855A, K810A and K848A mutants, the HNH communities are disposed in a circle and are connected by links, whose thickness is proportional to *Δ*EB (border sizes also correspond to the *Δ*EB of the specific community). Negative *Δ*EB (red) represent loss of communication, positive *Δ*EB (blue) indicate communication gain upon mutation.

The three K–*to*–A mutations also display important differences. Indeed, K855A mainly disrupts the communication at the level of the allosteric communities, while the non-allosteric sites gain in communication. The two other mutants, K810A and K848A, display a progressive gain in communication between both allosteric and non-allosteric sites. In these mutants, the allosteric A1 and A2 communities gain communication with the non-allosteric NA1 and NA2. This effect, which is more pronounced in K848A, reduces the negative impact of these mutations on the allosteric core of HNH. Analyses of the isolated HNH domain are consistent with these findings (Supplementary Fig. S9), confirming that K855A mainly disrupts the communication between sites that are highly involved in the allosteric pathway, while K810A and K848A also display gain in communication between allosteric and non-allosteric sites. In summary, our results show that the three K –*to*–A mutations alter the HNH allosteric structure, with K855A exerting a more pronounced perturbation of the signaling transfer. This links the specificity enhancement to alterations of the allosteric signaling. As we discuss below, the different alteration of the allosteric signaling observed here reflects the different capability of the three single mutants to enhance Cas9 specificity,^14^ offering important insights on the mechanistic basis of the specificity enhancement.

## DISCUSSION

Allostery is a fundamental property of the CRISPR-Cas9 gene editing tool.^32,33^ In this system, the allosteric relay is critical for transferring the information of DNA binding from the recognition (REC) lobe to the nuclease domains for cleavage and specificity.^4–8^ Biochemical,^4^ structural^9^ and biophysical^5–8^ approaches have shown that the HNH domain is the pivot of this allosteric regulation, holding a striking flexibility that facilitates the signal transduction^9,10^ and controls DNA cleavage.^4^

Here, we combined solution NMR, MD and network theory to reveal the allosteric role of three critical point mutations in HNH – K810A, K848A, and K855A – that increase the specificity of Cas9 and reduce its off-target activity.^14^ We first analyzed the possible structural perturbations induced by the presence of the specificity enhancement point mutations. The NMR Δδ perturbations revealed that K855A induces the most significant structural perturbations, while K810A and K848A progressively reveal weakened perturbations (Fig. 3). This was consistent with the computational assessment of the chemical environment, reporting a similar trend and a decreased EP score from K855A > K810A > K848A (Fig. 3C) and conveying that the three mutations differentially alter the HNH structure.

To characterize the allosteric signaling, we combined CPMG relaxation dispersion experiments with computational analyses based on graph theory that are suited for the detection of allosteric effects.^20–27^ We found that the three key mutations retain the overall dynamic pathway responsible for the information transfer, indicating that the allosteric signaling is preserved (Fig. 4 and Supplementary Figs. S3-5). We then employed community network analysis^26^ to identify the communication structure and how it is altered by the enhanced specificity mutations. In-depth analysis of the community network (Fig. 5) revealed that the three K–*to*–A mutations disrupt the main communication channel between RuvC and REC, as evidenced by a drop in the mutation-induced edge betweennesses difference (*Δ*EB) between the allosteric communities that are central in the communication pathway (Fig 5B). This indicates that an increase in specificity is associated to alterations of the HNH allosteric structure. Notably, K855A more strongly disrupts the communication between allosteric sites, compared to K848A and K810A that display a less negative impact (Fig. 5B). This is consistent with the structural perturbations observed through the NMR Δδ and the computed EP scores (Fig. 3), showing that the dynamical perturbation induced by K855A mutant is more pronounced than that generated by K810A and K848A variants. Interestingly, among the three HNH K–*to*–A mutations, K855A has shown to achieve the highest specificity enhancement as a single point mutation.^14^ Indeed, the specificity enhancement of the three single mutants toward the off-target validating VEGFA gene follows the K855A > K848A ~ K810A order.^14^ In the same study, K810A and K848A required further optimization to achieve maximal specificity. In light of these data, the specificity enhancement of the three single mutants reflects their different capability to differentially perturb the allosteric signaling.

On the basis of these observations, K855A, which is strongly impacts the HNH allosteric communication, could most prominently leverage the HNH allosterism to improve the system’s specificity. The HNH dynamics is indeed critical for the REC to RuvC signal transduction, and intervenes in the onset of off-target DNA cleavages, as shown by single-molecule and kinetic experiments,^5,6^ and by computational analysis.^8,12^ These studies observed remarkable alterations of the HNH dynamics upon binding of off-target DNA sequences at the level of the REC lobe. Hence, the alterations induced by K855A on the HNH structure and dynamics – which is the REC to RuvC allosteric pivot – are an indication that this mutant could increase the Cas9 specificity by modulating the HNH allosteric signaling and dynamical activation.

The K810A and K848A mutants, which affect the HNH structure and dynamics in a more moderate way, could exert a lower allosteric effect on the specificity enhancement. As noted above, these mutants require additional substitutions to achieve maximal specificity.^14^ The additional K1003A and R1060A locate remarkably away from HNH (Supplementary Fig. S10A) and might exploit different mechanisms to improve Cas9 specificity, such as altering the interactions with the distal bases of the DNA non-target strand.^14^ To understand further, we performed additional μs-length MD of the triple mutants K810A-K1003A-R1060A (*viz*., eSpCas9 1.0) and K848A-K1003A-R1060A (*viz*., eSpCas9 1.1), as well as of the WT Cas9 in an enlarged model system including a longer DNA non-target strand (Supplementary Fig. S10A, details in the Supplementary Data). As a result, K1003A and R1060A induce a remarkable flexibility of the distal DNA bases, compared to the WT Cas9 (Supplementary Fig. S10B). This is consistent with the hypothesis that the specificity enhancement of these mutants could also arise from the neutralization of positive charges interacting at the level of the DNA non-target strand.^14^ This could indeed favor the re-hybridization of DNA in the presence of off-target sequences, thereby limiting off-target cleavages.^34,35^

Taken together, these observations suggest that K855A could increase the specificity by mainly leveraging the HNH allosterism. Instead, K810A and K848A could combine more moderate allosteric effects with the weakening of the interactions at the level of the DNA non-target strand. This combination of allosteric and electrostatic effects could be critical for the triple mutants, which were optimized for both specificity and activity, with eSpCas9 1.1 being widely used in vitro.^14^ Our investigations thereby show that, in addition to electrostatic effects, the K–*to*–A mutants alter the dynamics of HNH, impacting the allosteric communication between REC and RuvC. ^4–8^ This altered communication has a profound influence on Cas9 activation, representing an important source for the observed specificity enhancement. In light of several mutational studies revealing an increase in function upon targeting allosteric sites (as reported, for instance, in several review articles),^20,36,37^ the critical hotspots and sites of synchronized dynamics identified here offer novel insights for mutational studies of CRISPR-Cas9, aimed at further ameliorating its function.

Finally, it is worth noting that the combination of solution NMR and molecular simulations enabled us to characterize the allosteric signaling from the individual HNH domain with experimental accuracy, to the full-length Cas9 with atomic level detail through MD simulations. This “bottom-up” approach exploits the capability of solution NMR to identify allosteric motions within optimized constructs of the multi-domain Cas9 protein,^11,28^ while all-atom MD simulations are used to characterize the communication network within the full-length Cas9. Future studies in our laboratories will leverage this approach to fully characterize the allosteric transmission across the multiple domains of Cas9 and its variants, gaining thorough insights on the system’s function and specificity.

## CONCLUSIONS

Here, the combination of solution NMR, molecular simulations and graph theory revealed that three lysine-to-alanine point mutations, which substantially increase the system’s specificity,^14^ alter the allosteric mechanism of information transfer in the CRISPR-Cas9 HNH endonuclease, impacting the signal transmission from the DNA recognition region to the catalytic sites. Specifically, the K855A mutation strongly disrupts the HNH domain allosteric structure, exerting the highest perturbation on the signaling transfer, while K810A and K848A result in more moderate effects on the allosteric intercommunication. This differential perturbation of the allosteric signaling reflects the different capability of the three single mutants to increase Cas9 specificity, and indicates that the allosteric communication is critical for the specificity enhancement. These findings are a step forward in the molecular level understanding of the CRISPR-Cas9 mechanism, and open the door for harnessing the allosteric signaling towards the improved system’s specificity.

## MATERIALS AND METHODS

### Protein Expression and Purification

The three enhanced specificity mutations (K810A, K848A, and K855A) of the *Streptococcus Pyogenes* Cas9 (SpCas9) protein were introduced into a previously reported construct of the HNH domain (residues 775-908) that has shown consistency with the structure of HNH within the full-length CRISPR-Cas9 system.^11 15^N labeled NMR samples were expressing by Rosetta(DE3) cells in M9 minimal containing MEM vitamins, MgSO_4_, and CaCl_2_. Cells were induced with 0.5 mM IPTG after reaching an OD600 of 0.8-0.9 and grown for 16-18 hours at 20°C post-induction. Cells were harvested by centrifugation and resuspended in 20 mM HEPES, 500 mM KCl, and 5 mM imidazole at pH 8.0. Cells were then lysed by ultrasonication and purified on Ni-NTA column with an elution buffer of 20 mM HEPES, 250 mM KCl, and 220 mM imidazole at pH 7.4. The N-terminal His-tag was removed by TEV protease. NMR samples were dialyzed into a buffer containing 20 mM HEPES, 80 mM KCl, 1 mM DTT, 1 mM EDTA, and 7.5% (v/v) D2O at pH 7.4.

### NMR Spectroscopy

NMR spin relaxation experiments were carried out at 600 and 850 MHz on Bruker Avance NEO and Avance III HD spectrometers, respectively. All NMR spectra were processed with NMRPipe^29^ and analyzed in NMRFAM-SPARKY.^38^ Carr-Purcell-Meiboom-Gill (CPMG)^30^ experiments were adapted from the report of Palmer and coworkers with a constant relaxation period of 40 ms and *v*_CPMG_ values of 25, 50×2, 100, 150, 200, 400, 500×2, 600, 800×2, 900, 1000 Hz. Relaxation dispersion curves were generated and exchange parameters were obtained from fits of these data carried out with RELAX^39^ using the R2eff, NoRex, Tollinger (TSMFK01), and Carver-Richards (CR72 and CR72-Full) models. Longitudinal and transfers relaxation rates were measured with relaxation times of 20×2, 60×2, 80, 200×2, 400, 800, 1000, and 1200 ms for T1 and 8.48, 16.96×2, 33.92, 67.84, 84.8×2, 101.76×2, 118.72, 135.68 where x2 represents repeated relaxation times. Peak intensities were quantified in SPARKY and the resulting exponential decays were fit in Mathematica. Steady-state ^1^H-[^15^N] Nuclear Overhouser Effect (NOE) were measured with a 6 second relaxation delay followed by a 3 second saturation (delay) for the saturated (unsaturated) experiments. All relaxation experiments were carried out in a temperature-compensated, interleaved manner. Model-free analysis using the Lipari-Szabo formalism^40^ was carried out on dual-field NMR data in RELAX with fully automated protocols.

### Molecular Dynamics (MD) simulations

MD simulations have been performed on the full-length CRISPR-Cas9 system and on the isolated form of the HNH domain. The X-ray structures of the full-length Sp CRISPR-Cas9 (5F9R.pdb, at 3.40 Å),^9^ and of the HNH construct (6O56.pdb, 1.90 Å resolution)^11^ have been used as models. Both systems have been considered as wild-type (WT) and introducing the three single point mutations K810A, K848A and K855A,^14^ resulting in eight simulation systems. All systems were solvated reaching periodic boxes of ~34,000 (isolated HNH) and ~340,000 (full-length Cas9) total atoms. A new AMBER ff99SBnmr2 force field,^41^ which improves the consistency of the backbone conformational ensemble with NMR experiments, was used for the protein. Nucleic acids were described including the ff99bsc0+χOL3 corrections for DNA^42^ and RNA.^43,44^ The TIP3P model was used for water.^45^ Simulations were performed in the NPT ensemble with temperature held at 310 K using the Bussi thermostat.^46^ The pressure was held at 1 bar with the Parrinello-Rahman barostat.^47^ A time step of 2 fs was applied (details are reported in the SI). A ~1.2 μs-long trajectory was collected in three replicas for the WT CRISPR-Cas9 system and for each of the K845A, K855A and H855A variants. The isolated HNH domain was also simulated in three replicas of ~1.2 μs each as WT and introducing the three K–*to*–A point mutations. This resulted in ~3.6 μs of MD for each simulated system and a total of ~14.4 μs of MD for the full-length CRISPR-Cas9 and also for the isolated HNH domain. This simulation length (in three replicates) was motivated by our previous work,^7,11^ showing that it provides a solid statistical ensemble for the purpose of the analysis of the allosteric mechanism (described below). Analysis of the results has been performed upon discarding the first ~200 ns of MD, to enable proper equilibration and a fair comparison. Data are reported for the separate replicas (Figs. S2, S4-S8), and for the overall ensemble (main text and Fig. S7-S9). Three additional systems, including a longer DNA non-target strand, were also built to simulate the triple mutants K810A-K1003A-R1060A (eSpCas9 1.0) and K848A-K1003A-R1060A (eSpCas9 1.1)^14^ and the WT Cas9 (details are in the Supplementary Information). These solvated systems comprised ~412,000 atoms and were also simulated for ~1.2 μs, each. All simulations were performed using Gromacs (v. 2018.3).^48^

### Environment Perturbation Score

To determine the structural perturbation induced by the three K–*to*–A mutations from MD simulations, we introduced an Environment Perturbation (EP) score measure. The EP score is a measure of the mutation-induced dynamic perturbation suffered by each residue in HNH given its local environment. The EP score analysis begins with the definition of a threshold radius *r_t_* around every heavy atom. A cutoff of 5 Å for *r_t_* has been used based on the typical upper distance between nuclei exhibiting a measurable NOE. Next, a frequency matrix *M* was created, whose elements *M_ij_* are the relative amount of time that residues *i* and *j* spend closer than *r_t_* during the MD simulation. Upon computing the matrix *M* for the WT system and for the K810A, K848A and K855A mutants, the EP score per residue, 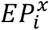, was calculated as follow:

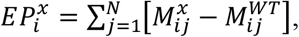

where *x* refers to the K810A, K848A or K855A mutants and *N* is the number of amino acids in HNH.

For every mutant *x*, the total strength of environmental perturbation (i.e., the total EP, 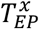) was defined as:

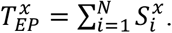

where:

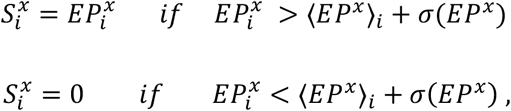

with the angular brackets representing the ensemble average and *σ*(*EP^x^*) the standard deviation along the simulations. According to the above, the EP score is a quantitative measure of the total heavy atom contact change for every residue, occurring as a consequence of the different mutations. This quantity thereby measures the overall change in chemical environment, and can be qualitatively compared with the composite NMR chemical shifts, which are a direct experimental reporter of the local environment.

### Communication Pathway Analysis

To describe the allosteric pathway of communication, information theory and network analysis have been applied to the analysis of our μs-length simulations.^26^ First, a Generalized Correlations (*GC*) analysis^49^ was carried out to compute the overall correlations between Cα atoms of the HNH domain. This analysis quantifies the system’s correlations based on the mutual information (*MI*)^49^ between two variables *x_i_* and *x_j_* (i.e., position vectors for the Cα atoms *i* and *j*):

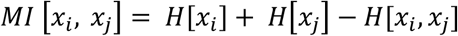

where *H* [*x_i_*] and *H* [*x_j_*] are the marginal Shannon entropies, and *H* [*x_i_*, *x_j_*] is the joint entropy, providing a link between motion correlations and information content (details are reported in the Supplementary Data). The *MI* can be converted into normalized *GC*, ranging from 0 (independent variables) to 1 (fully correlated variables):

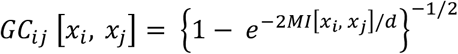

where *d* = 3 is the dimensionality of *x_i_* and *x_j_*. The *GC* values were used to build a network model, in which each residue was represented as a node connected by edges.^26^ The lengths of the edges were weighted using the *GC*, with the weight (*w_ij_*) of the edge connecting nodes *i* and *j* being:

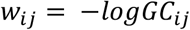

Hence, highly correlated pairs of residues are associated with efficient links for information transfer. Each node pair was connected by an edge if the residues involved spent at least 75 % of the simulation time within 5 Å. This threshold was carefully optimized in our previous studies.^7,11^

To determine the major channels of the information flow, we employed the optimal path search introduced by Dijkstra on our network models.^27^ The resulting pathways are composed by single-edge steps that maximize the total correlation (and optimize the momentum transport) between the signal “source” and “sink” amino acids. We studied the communication pathways that go through HNH from residues that connect REC (residues 789 and 794; sources) to RuvC (residues 841 and 858; sinks), and thereby transfer the information of DNA binding from the recognition region to the cleavage sites.^4–8^ The resulting routes maximize the dynamical cross-talk between the source and sink, serving as optimal communication channels. To account for the contribution of the most likely suboptimal pathways, the 10 shortest pathways for each source and sink pairs were computed, accumulated and plotted on the 3D structure of HNH in its isolated form and within the full-length Cas9.

### Community Network Analysis

To structure the allosteric network and how it is altered by the three K810A, K848A and K855A mutations, we performed Community Network Analysis.^26^ The dynamical network models described above were divided in local substructures – i.e., communities – composed of groups of nodes in which the network connections are dense but between which they are sparse. To structure the communities, the Girvan-Newman graph-partitioning approach^50^ was employed, using the Edge Betweenness (EB) as partitioning criterion (details are in the Supplementary Information). EB is defined as the number of shortest pathways that cross the edge, thereby accounting for the number of times an edge acts as a bridge in the communication flow between nodes of the network. The total EB between couples of communities (i.e., the sum of the EB of all edges connecting two communities) is a measure of their communication strength. Here, the total EB between couples of communities has been used to quantify the mutation-induced changes in the communication flowing through HNH. For each mutant (K855A, K810A and K484A), the mutation-induced EB change (*Δ*EB) has been computed as a difference between the EB of the mutant and the WT system. Normalized *Δ*EB range from negative (−1 < 0) to positive (0 < 1) values, indicating loss or gain in communication, respectively. Full details are reported in the Supplementary Information.

## ACKNOWLEDGMENTS

This material is based upon work supported by the National Institute of Health under Grant No. R01GM136815 (awarded to V.S.B., G.P. and G.P.L.) and Grant No. R01GM141329 (awarded to G.P). This work was also funded by the National Science Foundation under Grant No. CHE-1905374 (awarded to G.P.). Computer time for MD has been awarded by XSEDE under Grant No. TG-MCB160059 and by NERSC under Grant No. M3807 (to G.P.).

## AUTHOR CONTRIBUTIONS

L.N. performed molecular simulations and analyzed data. K.W.E. performed biochemical and NMR experiments. U.N.M. and P.R.A. analyzed data and developed analytical computational tools. V.S.B. overviewed research. G.P conceived and directed computational research. G.P.L. conceived and directed experimental research. G.P. wrote the manuscript with critical input from all authors.

## ADDITIONAL INFORMATION

Additional information is available as a supplementary information.

## DATA AVAILABILITY

The data that support the findings of this study are available from the corresponding author upon request.

## REFERENCES

1. Doudna, J. A. The promise and challenge of therapeutic genome editing. Nature 578, 229–236 (2020).

2. Jinek, M.; Chylinski, K.; Fonfara, I.; Hauer, M.; Doudna, J. A. & Charpentier, E. A Programmable Dual-RNA-Guided DNA Endonuclease in Adaptive Bacterial Immunity. Science 337, 816–821 (2012).

3. Jiang, F. & Doudna, J. A. CRISPR–Cas9 Structures and Mechanisms. Annu. Rev. Biophys. 46, 505–529 (2017).

4. Sternberg, S. H., LaFrance, B., Kaplan, M. & Doudna, J. A. Conformational control of DNA target cleavage by CRISPR–Cas9. Nature 527, 110–113 (2015).

5. Chen, J. S.; Dagdas, Y. S.; Kleinstiver, B. P.; Welch, M. M.; Sousa, A. A.; Harrington, L. B.; Sternberg, S. H.; Joung, J. K.; Yildiz, A. & Doudna, J. A. Enhanced Proofreading Governs CRISPR–Cas9 Targeting Accuracy. Nature 550, 407–410 (2017).

6. Dagdas, Y. S., Chen, J. S., Sternberg, S. H. & Doudna, J. A. A Conformational Checkpoint between DNA Binding and Cleavage by CRISPR-Cas9. Sci. Adv. 3, eaao002 (2017).

7. Palermo, G.; Ricci, C. G.; Fernando, A.; Basak, R.; Jinek, M.; Rivalta, I.; Batista, V. S. & McCammon, J. A. Protospacer Adjacent Motif-Induced Allostery Activates CRISPR-Cas9. J. Am. Chem. Soc. 139, 16028–16031 (2017).

8. Palermo, G.; Chen, J. S.; Ricci, C. G.; Rivalta, I.; Jinek, M.; Batista, V. S.; Doudna, J. A. & McCammon, J. A. Key Role of the REC Lobe during CRISPR–Cas9 Activation by ‘Sensing’, ‘Regulating’, and ‘Locking’ the Catalytic HNH Domain. Q. Rev. Biophys. 51, e9 (2018).

9. Jiang, F.; Taylor, D. W.; Chen, J. S.; Kornfeld, J. E.; Zhou, K.; Thompson, A. J.; Nogales, E. & Doudna, J. A. Structures of a CRISPR-Cas9 R-Loop Complex Primed for DNA Cleavage. Science 351, 867–871 (2016).

10. Palermo, G., Miao, Y., Walker, R. C., Jinek, M. & McCammon, J. A. Striking Plasticity of CRISPR-Cas9 and Key Role of Non-target DNA, as Revealed by Molecular Simulations. ACS Cent. Sci. 2, 756–763 (2016).

11. East, K. W.; Newton, J. C.; Morzan, U. N.; Narkhede, Y. B.; Acharya, A.; Skeens, E.; Jogl, G.; Batista, V. S.; Palermo, G. & Lisi, G. P. Allosteric Motions of the CRISPR-Cas9 HNH Nuclease Probed by NMR and Molecular Dynamics. J. Am. Chem. Soc. 142, 1348–1358 (2020).

12. Ricci, C. G.; Chen, J. S.; Miao, Y.; Jinek, M.; Doudna, J. A.; McCammon, J. A. & Palermo, G. Deciphering Off-Target Effects in CRISPR-Cas9 through Accelerated Molecular Dynamics. ACS Cent. Sci. 5, 651–662 (2019).

13. Kleinstiver, B. P.; Pattanayak, V.; Prew, M. S.; Tsai, S. Q.; Nguyen, N. T.; Zheng, Z. & Joung, J. K. High-Fidelity CRISPR–Cas9 Nucleases with No Detectable Genome-Wide off-Target Effects. Nature 529, 490–495 (2016).

14. Slaymaker, I. M.; Gao, L.; Zetsche, B.; Scott, D. A.; Yan, W. X. & Zhang, F. Rationally Engineered Cas9 Nucleases with Improved Specificity. Science 351, 84–88 (2016).

15. Casini, A.; Olivieri, M.; Petris, G.; Montagna, C.; Reginato, G.; Maule, G.; Lorenzin, F.; Prandi, D.; Romanel, A.; Demichelis, F.; Inga, A. & Cereseto, A. A Highly Specific SpCas9 Variant Is Identified by in Vivo Screening in Yeast. Nat. Biotechnol. 36, 265–271 (2018).

16. Tzeng, S. R. & Kalodimos, C. G. Protein dynamics and allostery: An NMR view. Current Opin. Struct. Biol. 21 62–67 (2011).

17. Lisi, G. P. & Loria, J. P. Solution NMR Spectroscopy for the Study of Enzyme Allostery. Chem. Rev. 116, 6323–6369 (2016).

18. Grutsch, S., Brüschweiler, S. & Tollinger, M. NMR Methods to Study Dynamic Allostery. PLOS Comput. Biol. 12, e1004620 (2016).

19. Boulton, S. & Melacini, G. Advances in NMR Methods to Map Allosteric Sites: From Models to Translation. Chem. Rev. 116, 6267–6304 (2016).

20. Liu, J. & Nussinov, R. Allostery: An Overview of Its History, Concepts, Methods, and Applications. PLOS Comput. Biol. 12, e1004966 (2016).

21. Guo, J. & Zhou, H. X. Protein Allostery and Conformational Dynamics. Chem. Rev. 116, 6503–6515 (2016).

22. Dokholyan, N. V. Controlling Allosteric Networks in Proteins. Chem. Rev. 116, 6463–6487 (2016).

23. East, K. W.; Skeens, E.; Cui, J. Y.; Belato, H. B.; Mitchell, B.; Hsu, R.; Batista, V. S.; Palermo, G. & Lisi, G. P. NMR and Computational Methods for Molecular Resolution of Allosteric Pathways in Enzyme Complexes. Biophys. Rev. 12, 155–174 (2020).

24. Vendruscolo, M. The statistical theory of allostery. Nat. Chem. Biol. 7, 411–412 (2011).

25. Wodak, S. J.; Paci, E.; Dokholyan, N. V.; Berezovsky, I. N.; Horovitz, A.; Li, J.; Hilser, V. J.; Bahar, I.; Karanicolas, J.; Stock, G.; Hamm, P.; Stote, R. H.; Eberhardt, J.; Chebaro, Y.; Dejaegere, A.; Cecchini, M.; Changeux, J.-P.; Bolhuis, P. G.; Vreede, J.; Faccioli, P.; Orioli, S.; Ravasio, R.; Yan, L.; Brito, C.; Wyart, M.; Gkeka, P.; Rivalta, I.; Palermo, G.; McCammon, J. A.; Panecka-Hofman, J.; Wade, R. C.; Di Pizio, A.; Niv, M. Y.; Nussinov, R.; Tsai, C.-J.; Jang, H.; Padhorny, D.; Kozakov, D. & McLeish, T. Allostery in Its Many Disguises: From Theory to Applications. Structure 27, 566–578 (2019).

26. Sethi, A., Eargle, J., Black, A. A. & Luthey-Schulten, Z. Dynamical networks in tRNA: protein complexes. Proc. Natl. Acad. Sci. U. S. A. 106, 6620–6625 (2009).

27. Bowerman, S. & Wereszczynski, J. Detecting Allosteric Networks Using Molecular Dynamics Simulation. Methods Enzymol. 578, 429–447 (2016).

28. Nerli, S., De Paula, V. S., McShan, A. C. & Sgourakis, N. G. Backbone-independent NMR resonance assignments of methyl probes in large proteins. Nat. Commun. 12, 1–13 (2021).

29. Delaglio, F.; Grzesiek, S.; Vuister, G. W.; Zhu, G.; Pfeifer, J. & Bax, A. NMRPipe: A Multidimensional Spectral Processing System Based on UNIX Pipes. J. Biomol. NMR 6, 277–293 (1995).

30. Loria, J. P., Rance, M. & Palmer 3rd, A. G. A Relaxation-Compensated Carr-Purcell-Meiboom-Gill Sequence for Characterizing Chemical Exchange by NMR Spectroscopy. J. Am. Chem. Soc. 121, 2331–2332 (1999).

31. Oyen, D.; Bryn Fenwick, R.; Aoto, P. C.; Stanfield, R. L.; Wilson, I. A.; Jane Dyson, H. & Wright, P. E. Defining the Structural Basis for Allosteric Product Release from E. Coli Dihydrofolate Reductase Using NMR Relaxation Dispersion. J. Am. Chem. Soc. 139, 11233–11240 (2017).

32. Nierzwicki, Ł., Arantes, P. R., Saha, A. & Palermo, G. Establishing the Allosteric Mecanism in CRISPR-Cas9. WIREs Comput. Mol. Sci. 11, e1503, (2021).

33. Zuo, Z. & Liu, J. Allosteric regulation of CRISPR-Cas9 for DNA-targeting and cleavage. Curr. Opin. Struct. Biol. 62, 166–174 (2020).

34. Singh, D., Sternberg, S. H., Fei, J., Doudna, J. A. & Ha, T. Real-time observation of DNA recognition and rejection by the RNA-guided endonuclease Cas9. Nat Commun 7, 12778 (2016).

35. Singh, D.; Wang, Y.; Mallon, J.; Yang, O.; Fei, J.; Poddar, A.; Ceylan, D.; Bailey, S. & Ha, T. Mechanisms of Improved Specificity of Engineered Cas9s Revealed by Single-Molecule FRET Analysis. Nat. Struct. Mol. Biol. 25, 347–354 (2018).

36. Guo, J. & Zhou, H. X. Protein Allostery and Conformational Dynamics. Chem. Rev. 116, 6503–6515 (2016).

37. Dokholyan, N. V. Controlling Allosteric Networks in Proteins. Chem. Rev. 116, 6463–6487 (2016).

38. Lee, W., Tonelli, M. & Markley, J. L. NMRFAM-SPARKY: Enhanced software for biomolecular NMR spectroscopy. Bioinformatics 31, 1325–1327 (2015).

39. Bieri, M., d’Auvergne, E. J. & Gooley, P. R. relaxGUI: A New Software for Fast and Simple NMR Relaxation Data Analysis and Calculation of ps-ns and us Motion of Proteins. J. Biomol. NMR 50, 147–155 (2011).

40. Lipari, G. & Szabo, A. Model-Free Approach to the Interpretation of Nuclear Magnetic Resonance Relaxation in Macromolecules. 1. Theory and Range of Validity. J. Am. Chem. Soc. 104, 4546–4559 (1982).

41. Yu, L., Li, D. W. & Brüschweiler, R. Balanced Amino-Acid-Specific Molecular Dynamics Force Field for the Realistic Simulation of Both Folded and Disordered Proteins. J. Chem. Theory Comput. 16, 1311–1318 (2020).

42. Perez, A.; Marchan, I.; Svozil, D.; Sponer, J.; Cheatham, T. E. 3rd; Laughton, C. A. & Orozco, M. Refinement of the AMBER Force Field for Nucleic Acids: Improving the Description of Alpha/Gamma Conformers. Biophys. J. 92, 3817–3829 (2007).

43. Zgarbova, M.; Otyepka, M.; Sponer, J.; Mladek, A.; Banas, P.; Cheatham, T. E. & Jurecka, P. Refinement of the Cornell et Al. Nucleic Acids Force Field Based on Reference Quantum Chemical Calculations of Glycosidic Torsion Profiles. J. Chem. Theory Comput. 7, 2886–2902 (2011).

44. Banas, P.; Hollas, D.; Zgarbova, M.; Jurecka, P.; Orozco, M.; Cheatham, T. E. 3rd; Sponer, J. & Otyepka, M. Performance of Molecular Mechanics Force Fields for RNA Simulations: Stability of UUCG and GNRA Hairpins. J. Chem. Theor. Comput. 6, 3836–3849 (2010).

45. Jorgensen, W. L., Chandrasekhar, J., Madura, J. D., Impey, R. W. & Klein, M. L. Comparison of simple potential functions for simulating liquid water. J. Chem. Phys. 79, 926–935 (1983).

46. Bussi, G., Donadio, D. & Parrinello, M. Canonical sampling through velocity rescaling. J. Chem. Phys. 126, 014101 (2007).

47. Parrinello, M. & Rahman, A. Polymorphic transitions in single crystals: A new molecular dynamics method. J. Appl. Phys. 52, 7182–7190 (1981).

48. Van Der Spoel, D.; Lindahl, E.; Hess, B.; Groenhof, G.; Mark, A. E. & Berendsen, H. J. C. GROMACS: Fast, Flexible, and Free. J. Comput. Chem. 26, 1701–1718 (2005).

49. Lange, O. F. & Grubmüller, H. Generalized correlation for biomolecular dynamics. Proteins Struct. Funct. Genet. 62, 1053–1061 (2006).

50. Girvan, M. & Newman, M. E. J. Community structure in social and biological networks. Proc. Natl. Acad. Sci. U. S. A. 99, 7821–6 (2002).

